# Dietary Sodium Restriction Reprograms Gut Microbial Fermentation and Reduces Host Energy Harvest

**DOI:** 10.64898/2026.04.20.719706

**Authors:** Joshua Cornman-Homonoff, Kumaran M. Rajendran, Saravanan Kolandaivelu, Steven D. Coon, Justin T. Kupec, Lei Wang, Gangqing Hu, Venkatakrishna R. Jala, Geoffrey I. Sandle, Vazhaikkurichi M. Rajendran

## Abstract

Diet is a major determinant of gut microbiome structure and function, yet the role of dietary electrolytes—particularly sodium—remains poorly defined. Here, we identify dietary sodium availability as a key regulator of gut microbial fermentation and host energy harvest. Using a controlled sodium-sufficient versus sodium-deprived dietary intervention in rats, we integrated shotgun metagenomic sequencing, functional pathway analysis, targeted short-chain fatty acid (SCFA) quantification, and host physiological phenotyping. Sodium deprivation induced a coordinated restructuring of the gut microbiome, characterized by depletion of classical saccharolytic Firmicutes, including multiple *Lactobacillus* species, and enrichment of stress-tolerant, metabolically flexible taxa. Functional profiling revealed a shift away from growth-associated metabolic programs toward stress-adaptive and nutrient-scavenging pathways. Consistent with these changes, fecal concentrations of key SCFAs—including acetate, butyrate, hexanoate, and valerate—were significantly reduced, indicating impaired microbial fermentative capacity. These microbiome-level alterations translated into measurable host phenotypes, including reduced cecal mass and attenuated weight gain, consistent with decreased microbial energy harvest. Together, these findings establish a functional link between luminal sodium availability, microbial metabolic efficiency, and host energy balance, extending the framework of diet–microbiome interactions beyond macronutrients to include dietary electrolytes. This work identifies sodium as a previously underappreciated ecological constraint shaping gut microbial metabolism and suggests that modulation of dietary sodium intake may influence host metabolic outcomes through microbiome-mediated mechanisms.

## INTRODUCTION

The human gastrointestinal tract harbors a complex and dynamic community of trillions of microorganisms, collectively referred to as the gut microbiota (1). This includes bacteria, viruses, and other microbes, with bacteria being the most abundant and extensively studied. The predominant bacterial phyla in the gut microbiota are Firmicutes, Bacteroidetes, Proteobacteria, Actinobacteria, and Verrucomicrobia (2). Among these, Firmicutes (or Bacillota; Gram-positive) and Bacteroidetes (or Bacteroidota; Gram-negative) are the most abundant (3, 4). Although microbes are distributed throughout the gastrointestinal tract, the highest density (approximately 10¹¹ per gram feces) is found in the distal colon (5). A well-balanced gut microbiota is essential for maintaining host health, playing critical roles in metabolism, immune function, and intestinal homeostasis. Conversely, dysbiosis, characterized by a reduction or alteration in microbial diversity, has been associated with a range of metabolic and inflammatory disorders, including obesity, diabetes, and inflammatory bowel diseases (6–8). Among various environmental factors, diet is one of the most influential in shaping the composition and function of the gut microbiota (9). As such, there is increasing interest in modulating the gut microbiota through dietary and nutritional interventions to promote gut health and serve as adjunctive strategies for treating obesity and related diseases (10–14).

Although salt (NaCl) is an essential component of human diet, excess Na^+^ consumption is associated with hypertension (15, 16), while a reduced Na^+^ intake lowers blood pressure (17, 18). Recent studies have shown significant alterations in gut microbiota to be associated with the development of hypertension in both humans and animals (19–23). Given the association between dietary Na^+^ intake and hypertension, and the contact between ingested Na^+^ and the gastrointestinal tract, there are likely to be important interactions between dietary Na^+^ and the gut microbiota. High dietary Na^+^ intakes have been shown to have a negative impact on gut microbial composition, which leads to inflammation and hypertension (23–25), whereas low Na^+^ intakes have inconsistent effects on microbial composition (26). The present study was designed to determine whether dietary Na^+^ deprivation alters fecal microbial composition in rats. We found that Na^+^ deprivation significantly changed the microbiota profile, decreased fecal concentrations of short chain fatty acids (SCFAs), and slowed the rate of body weight gain, raising the possibility that dietary Na^+^ deprivation may be a useful additional strategy in the management of obesity.

## METHODS

### Animals and fecal collection

Male Sprague Dawley rats (125 – 150 g; Charles River, Raleigh, NC) housed 2 rats/cage were fed either a Na^+^-sufficient (NaS) diet or a Na^+^-deprived (NaD; cat# MP296023210; MP Biomedicals) diet, and water ad libitum for 7 days. Starting from day-1, animals were weighed daily to determine weight gain. Food and water (ml) consumptions were determined per cage. On day-8, rats were anaesthetized with 5% isoflurane and euthanized with high dose isoflurane/cervical dislocation. Blood was collected by cardiac puncture, while the entire colon and caecum were removed. Following photography and length measurement, 2-3 fecal pellets were extracted from the colon, placed in sterile microfuge tubes, and stored -80°C. Distal colons flushed with ice-cold phosphate buffered saline (PBS) were used for physiological, histological, and immunofluorescence studies. Fecal samples were shipped on dry ice to Microbiome Insights Inc, Crestwood Place, Richmond, BC, Canada for microbiome and SCFA analyses. All experimental procedures used in this study were approved by West Virginia University’s Institutional Animal Care and Use Committee (WVU-IACUC).

### Shotgun metagenomic sequencing and quality control

Fecal DNA was extracted by Microbiome Insights (Richmond, BC, Canada) using the Qiagen MagAttract PowerSoil DNA KF kit (formerly MO BIO PowerSoil DNA Kit) on a KingFisher automated platform. DNA quality was assessed by agarose gel electrophoresis and quantified using a Qubit 3.0 fluorometer (Thermo Fisher Scientific). Sequencing libraries were prepared using the Illumina Nextera DNA library preparation kit following the manufacturer’s protocol. Shotgun metagenomic sequencing was performed on an Illumina NextSeq platform using 2 × 150 bp paired-end reads. Demultiplexed reads underwent initial quality assessment using FastQC v0.11.5. Quality control included paired-end read joining using FLASH v1.2.11, removal of host-derived sequences by alignment to the human reference genome (hg19 / GRCh37), and quality trimming using Trimmomatic v0.36 with custom parameters (LEADING:5, TRAILING:5, SLIDINGWINDOW:4:15, MINLEN:70). Following quality control, a median of approximately 13.4 million high-quality reads per sample was retained for downstream analyses.

### Taxonomic Profiling

#### Taxonomic classification

Taxonomic composition was determined from quality-filtered reads using MetaPhlAn2, which profiles microbial community composition based on clade-specific marker genes. Relative abundances were summarized at multiple taxonomic levels, including phylum, class, order, genus, and species. Community composition plots were generated using normalized relative abundance values.

### Functional Profiling

#### Functional annotation and pathway analysis

Functional profiling was performed by mapping quality-filtered reads to functional genes and summarizing them into pathways using MetaCyc pathway definitions. Approximately 50% of reads were assigned to annotated functional genes. Pathway richness (number of unique pathways per sample) was calculated, and differences between dietary groups were assessed using non-parametric statistical testing (see Statistics section below)

### Fecal short-chain fatty acid quantification

Fecal short-chain fatty acids (SCFAs) were quantified by Microbiome Insights using gas chromatography with flame ionization detection (GC-FID). Frozen fecal samples were thawed on ice and homogenized in acidified aqueous solution to extract SCFAs. Samples were acidified to protonate fatty acids and centrifuged to remove particulate matter. The clarified supernatant was transferred to GC vials for analysis. SCFA separation was performed on a capillary GC column optimized for volatile fatty acids, and detection was achieved using a flame ionization detector. Individual SCFAs were identified by comparison of retention times to authenticated external standards and quantified using calibration curves generated from known concentrations of SCFA standards. An internal standard was included to correct for extraction and injection variability. The SCFAs quantified included acetate, propionate, butyrate, isobutyrate, valerate, isovalerate, and hexanoate. Concentrations were reported as absolute abundances normalized to fecal sample mass. Quality control procedures included assessment of chromatographic peak resolution, internal standard recovery, and calibration linearity across the analytical range.

### Statistical analysis

All statistical analyses were performed on samples with n = 10 animals per group. Alpha diversity metrics (e.g., Shannon diversity) and pathway richness were compared between dietary groups using non-parametric tests, including the Kruskal–Wallis test, where appropriate. Beta diversity analyses were conducted using Bray–Curtis dissimilarity matrices, which account for both presence/absence and relative abundance of taxa or pathways. Ordination was performed using non-metric multidimensional scaling (NMDS) to visualize differences in microbial community structure and functional profiles between dietary groups. Group-level differences in overall taxonomic and functional composition were evaluated using permutational multivariate analysis of variance (PERMANOVA) implemented via the adonis function in the vegan R package. The coefficient of determination (R²) was used to estimate the proportion of variance explained by dietary treatment, with statistical significance assessed by permutation testing. Differential abundance testing of individual taxa and functional pathways was performed using DESeq2. P-values were adjusted for multiple testing using the Benjamini–Hochberg false discovery rate (FDR) method. Adjusted P values < 0.05 were considered statistically significant unless otherwise specified. SCFAs statistical analyses were performed using n = 10 animals per group. SCFA concentrations were assessed for normality prior to hypothesis testing. Because several SCFA distributions deviated from normality, non-parametric statistical tests were applied. Comparisons between NaS and NaD groups were conducted using the Mann–Whitney U test for individual SCFAs. Results are presented as individual biological replicates with group summaries. A two-sided P value < 0.05 was considered statistically significant. No imputation was performed for values below the limit of detection; these were retained as measured. Statistical analyses were conducted using standard statistical software.

## RESULTS

### Dietary Na^+^-deprivation alters gut microbial community structure

Na^+^-deprivation resulted in significant restructuring of overall gut microbial community composition. Alpha diversity, assessed using the Shannon index, was significantly higher in NaD rats compared NaS controls, indicating increased within-sample taxonomic diversity under low-Na^+^ conditions (Fig. 1A). Community-level differences were evaluated using Bray–Curtis dissimilarity followed by permutational multivariate analysis of variance (PERMANOVA). Dietary Na^+^ status explained a significant proportion of variance in microbial composition (PERMANOVA, P < 0.01), demonstrating that Na^+^ availability is a major determinant of gut microbiome structure (Fig. 1B and Supplementary Table-ST1). Ordination analysis revealed clear separation between NaS and NaD samples, consistent with reproducible Na^+^-dependent community restructuring. At the phylum level, differential abundance analysis using DESeq2 identified significant depletion of Firmicutes and enrichment of Bacteroidetes, Actinobacteria, and Proteobacteria in NaD rats relative to NaS controls (Fig. 1C–I and Supplementary Table-ST2; FDR-adjusted P < 0.05). These shifts resulted in a significantly reduced Firmicutes:Bacteroidetes ratio in NaD animals, reflecting a coordinated redistribution of dominant gut microbial phyla under Na^+^-restricted conditions.

**Figure 1.**
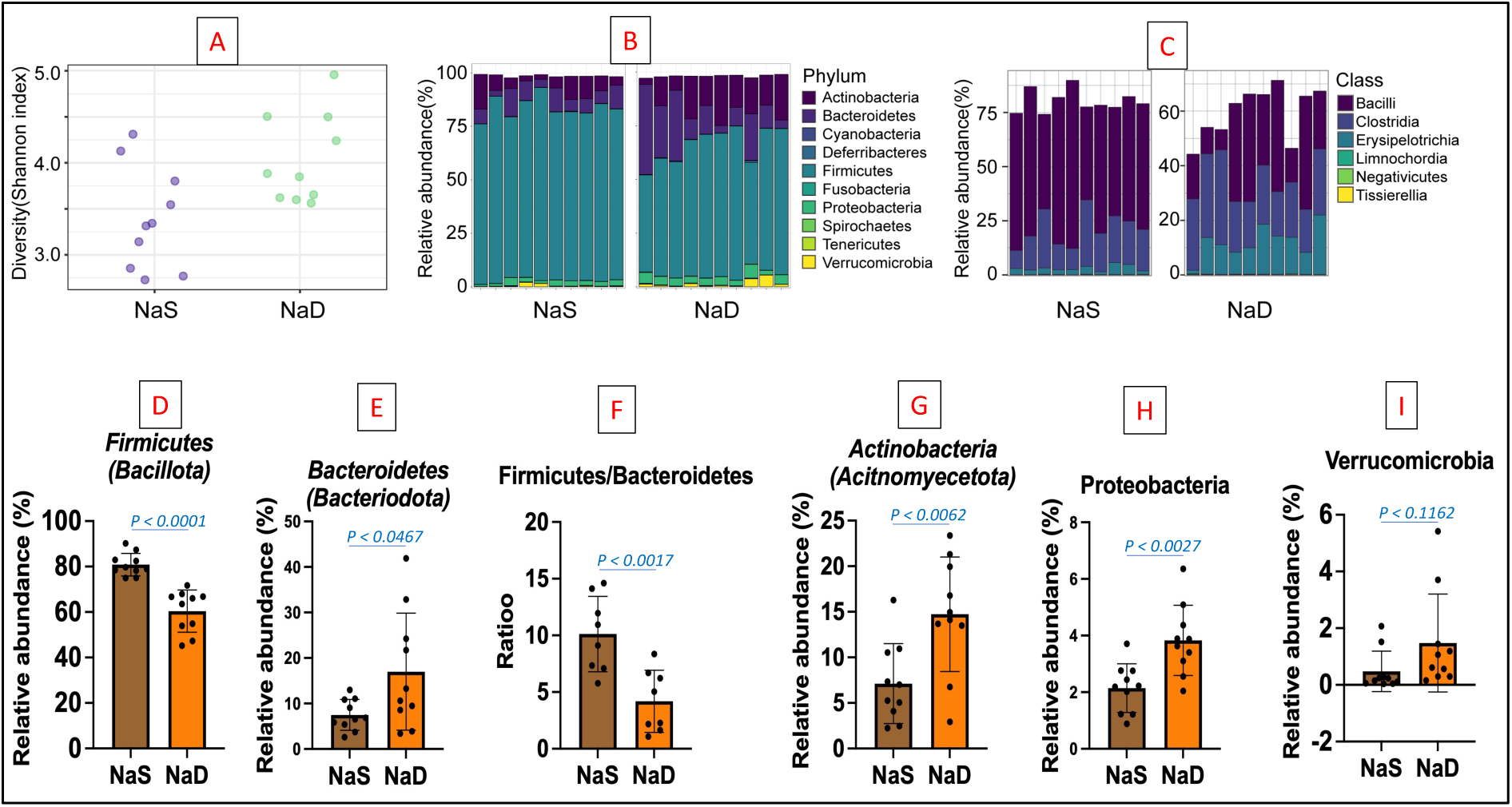
Dietary Na^+^-deprivation restructures gut microbial community composition. (A) Shannon alpha-diversity indices showing increased within-sample diversity in Na^+^-deprived (NaD) rats compared with Na^+^-sufficient (NaS) controls. (B) Bray–Curtis dissimilarity ordination demonstrating significant separation of microbial communities by dietary Na^+^ status (PERMANOVA). (C–I) Differential abundance of major bacterial phyla and Firmicutes:Bacteroidetes ratio assessed by DESeq2. NaD rats exhibited significant depletion of Firmicutes and enrichment of Bacteroidetes, Actinobacteria, and Proteobacteria (FDR-adjusted P < 0.05). Boxes represent interquartile range with median; whiskers denote range; points represent biological replicates.

### Na^+^-deprivation drives species-specific restructuring within Lactobacillaceae

To identify Na^+^-sensitive taxa contributing to community-level changes, species-level differential abundance analysis was performed using DESeq2 on normalized shotgun metagenomic counts. Multiple Lactobacillus and Limosilactobacillus species exhibited significant Na^+^-dependent abundance changes (Fig. 2 and Supplementary Table-ST2). DESeq2 analysis revealed significant depletion of Limosilactobacillus fermentum, Lactobacillus amylolyticus, Lactobacillus reuteri in NaD rats compared with NaS controls (FDR-adjusted P < 0.01). In contrast, Limosilactobacillus vaginalis showed relative enrichment under Na^+^-deprived conditions, indicating divergent species-specific responses within Lactobacillaceae. These results demonstrate that Na^+^-deprivation does not uniformly suppress Lactobacillus-related taxa, but instead selectively reshapes species composition, favoring taxa with greater osmo-adaptive or niche-specific metabolic capacity while depleting classical saccharolytic members.

**Figure 2.**
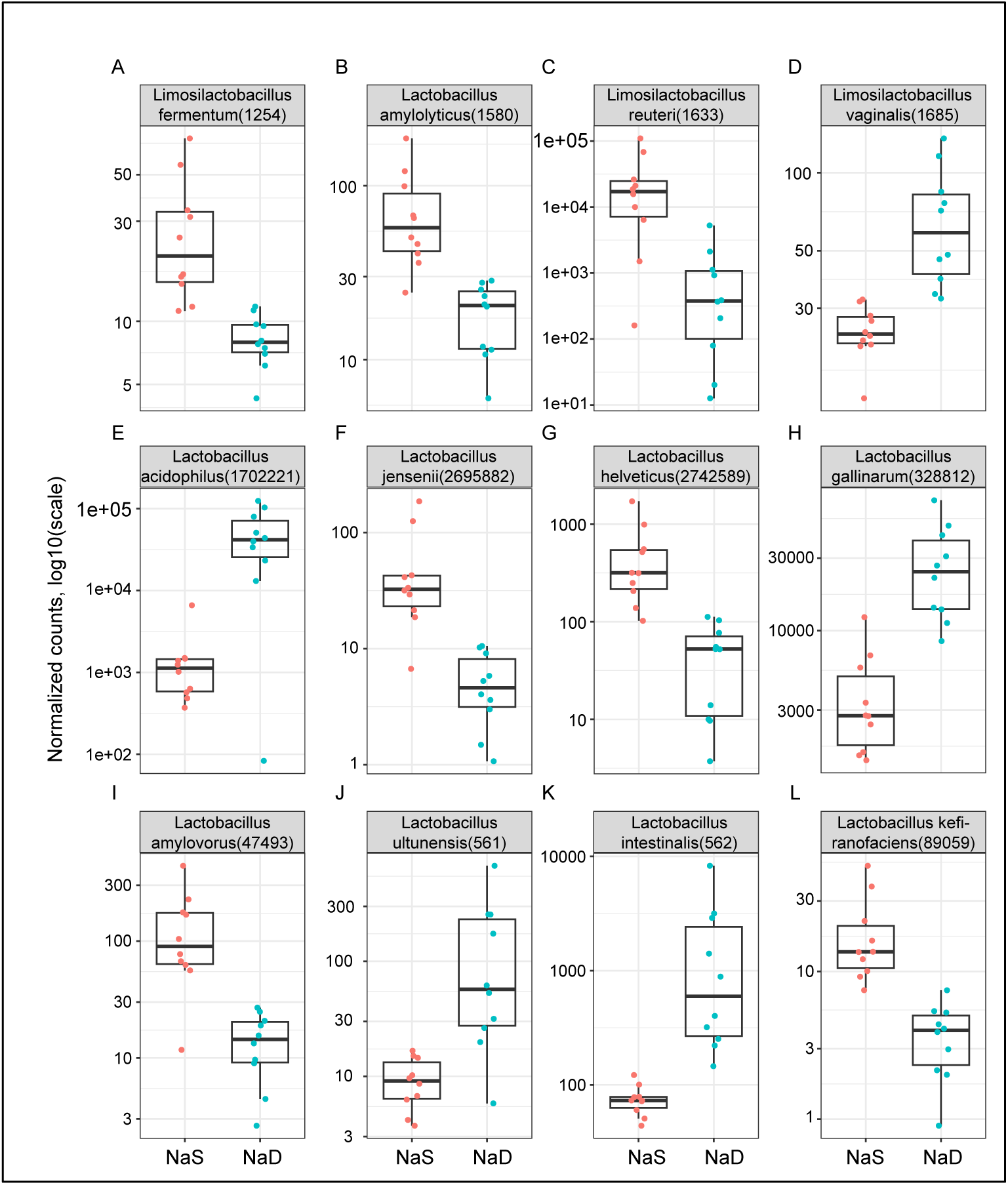
Na^+^-deprivation induces species-specific restructuring within Lactobacillaceae. (A–F) DESeq2-based differential abundance analysis of selected Lactobacillus and Limosilactobacillus species using normalized shotgun metagenomic read counts (log₁₀ scale). Several saccharolytic species (L. iners, L. apis, L. crispatus, L. fermentum, L. reuteri) were significantly depleted in Na^+^-deprived (NaD) rats, whereas Limosilactobacillus vaginalis was relatively enriched (FDR-adjusted P < 0.01). Boxes indicate interquartile range with median; whiskers denote range; points represent biological replicates.

### Dietary Na^+^ restriction reprograms microbial functional profiles

Functional pathway composition was assessed using Bray–Curtis dissimilarity followed by PERMANOVA. Dietary Na^+^ status significantly influenced overall functional pathway structure (PERMANOVA, P < 0.05), indicating coordinated Na^+^-dependent shifts in microbial metabolic potential (Supplementary Figs. S1–S20). Inspection of pathway abundance profiles revealed consistent reductions in pathways associated with microbial growth and fermentative metabolism, including carbohydrate metabolism, amino-acid metabolism, lipid and membrane biosynthesis, ribosomal functions, and electron-transport–related processes in NaD microbiomes (Supplementary Figs. S1A–S1M). Conversely, NaD microbiomes exhibited relative enrichment of pathways associated with environmental stress response, nutrient scavenging, iron acquisition, RNA modification, and secretion systems, including siderophore biosynthesis and two-partner secretion pathways (Supplementary Figs. S1O–S1T). Although no individual functional pathways reached statistical significance after multiple-testing correction, the consistent directional shifts across pathway groups, together with significant PERMANOVA results, indicate coordinated Na⁺-dependent functional reprogramming of the gut microbiome. Global functional profiling at SEED Subsystems Levels 1 and 2 demonstrated broad restructuring of microbial functional capacity under NaD (Supplementary Figs. S2 and S3), consistent with pathway-level changes identified by differential abundance analysis.

### Dietary Na^+^-deprivation reduces microbial short-chain fatty acid output

Consistent with Na^+^-dependent suppression of microbial fermentative activity, fecal short-chain fatty acid (SCFA) profiles differed between NaS and NaD) rats (Fig. 3 and Supplementary Table-ST3). NaD animals exhibited robust reductions in the major fermentative SCFAs acetate, butyrate, and hexanoate, with acetate significantly decreased (P = 0.002) and butyrate and hexanoate showing the largest magnitude reductions (P < 0.001 for both). Valerate concentrations were modestly but significantly reduced in NaD rats (P = 0.0350), indicating that sodium restriction also affects select secondary fermentation products. In contrast, propionate displayed a non-significant downward trend (P = 0.063), while isobutyrate (P = 0.116) and isovalerate (P = 0.284) did not differ significantly between groups. Together, these data indicate that dietary sodium restriction preferentially suppresses key microbial fermentation outputs that contribute substantially to host energy harvest, while exerting more limited effects on branched-chain SCFAs derived primarily from amino-acid fermentation.

**Figure 3.**
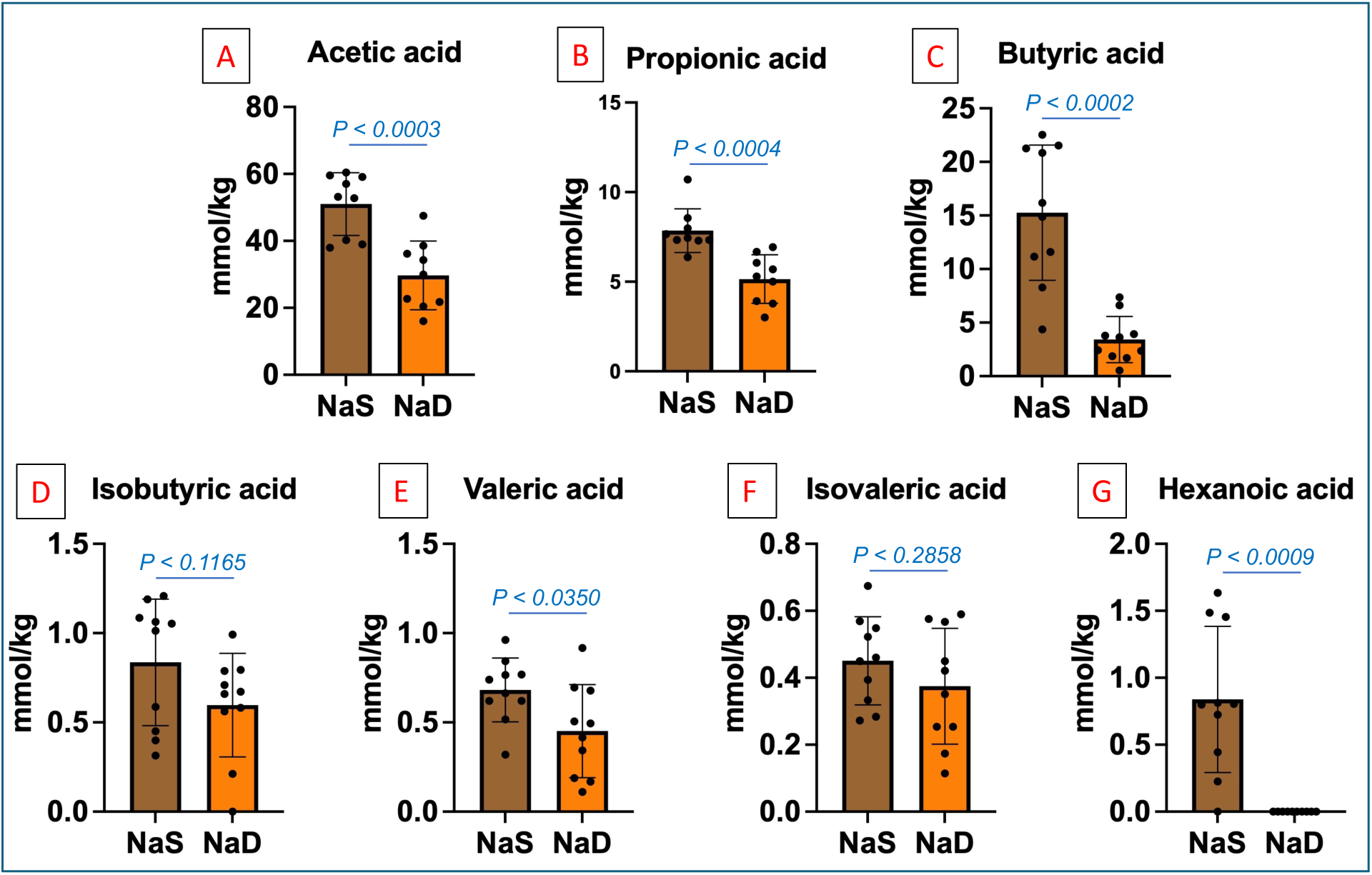
Dietary Na^+^ restriction reduces fecal short-chain fatty acid concentrations. (A–G) Fecal concentrations of acetate, propionate, butyrate, isobutyrate, valerate, isovalerate, and hexanoate in Na^+^-sufficient (NaS) and Na^+^-deprived (NaD) rats (n = 10 per group). Acetate concentrations were significantly reduced in NaD animals (P = 0.002), while butyrate and hexanoate exhibited pronounced reductions (P < 0.001 for both). Propionate showed a non-significant trend toward reduction (P = 0.063). Valerate concentrations were significantly reduced in NaD animals (P = 0.0350), whereas propionate, isobutyrate (P = 0.116), and isovalerate (P = 0.284) did not differ significantly between groups. Group comparisons were performed using two-sided Mann–Whitney U tests.

### Reduced microbial energy harvest is associated with altered host physiology

To assess whether Na^+^-dependent microbiome remodeling translated into host phenotypic changes, body weight, intake behavior, and gut morphology were evaluated. NaD rats exhibited reduced body weight gain over the experimental period compared with NaS controls (Fig. 4A–B). Although modest reductions in food and water intake were observed, these differences were insufficient to fully explain the attenuated weight gain. NaD rats also exhibited a marked reduction in cecal size and mass (Fig. 4E–G), a classic indicator of diminished microbial fermentation and substrate turnover, while colon length remained unchanged. These findings support a model in which dietary Na^+^ restriction limits host energy harvest primarily through microbiome-mediated suppression of fermentative capacity rather than through structural changes to the intestine.

**Figure 4.**
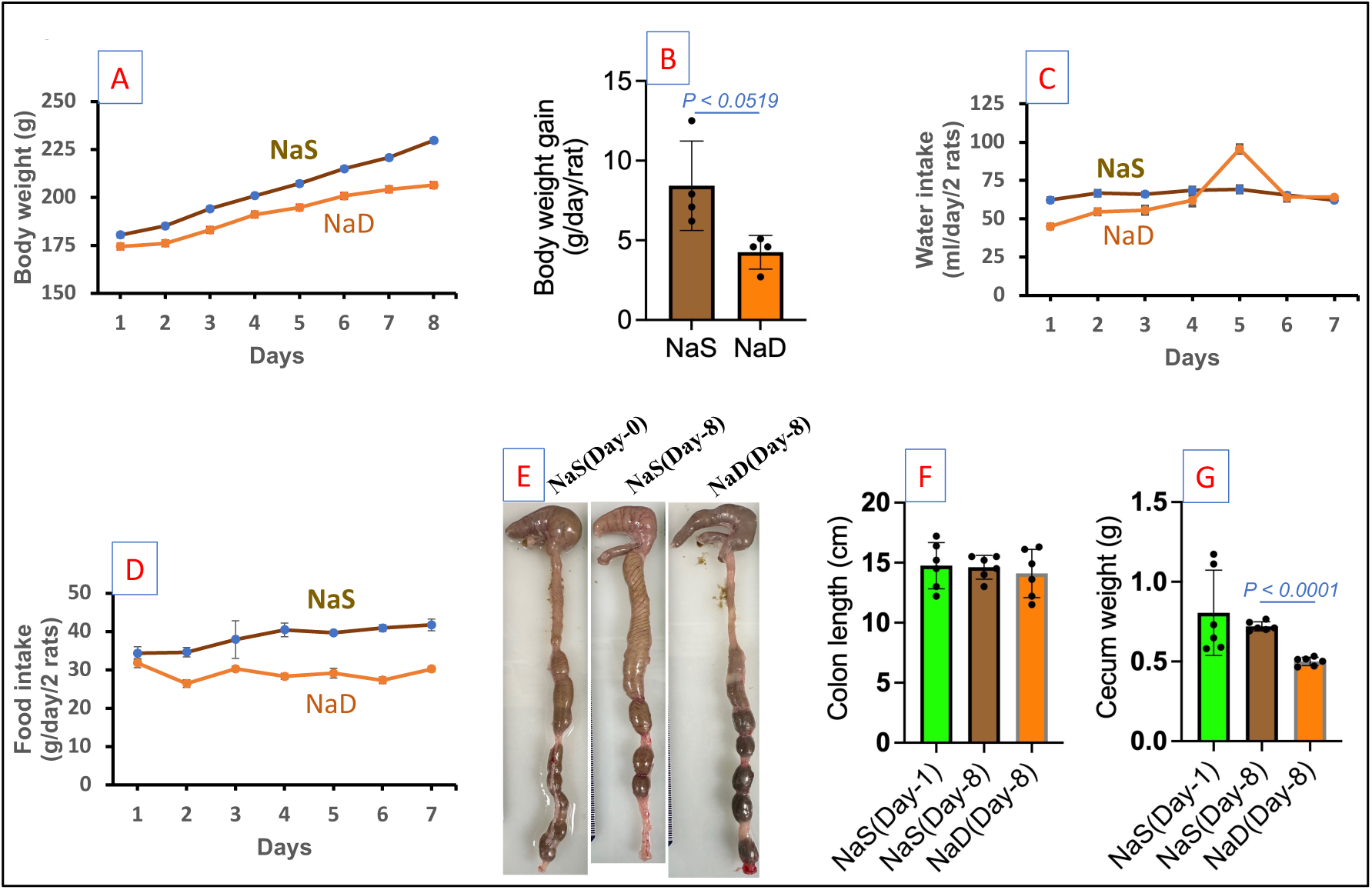
Na^+^-deprivation attenuates weight gain and reduces cecal mass. (A–B) Body-weight trajectories and mean daily weight gain in NaS and NaD rats. (C–D) Daily water and food intake. (E–G) Cecal size, colon length, and cecal weight. NaD rats exhibited significantly reduced cecal mass without changes in colon length, consistent with reduced microbial fermentative activity.

## DISCUSSION

Dietary Na^+^ restriction is widely recommended for the prevention and management of hypertension and cardiometabolic disease; however, its effects on gut microbial ecology and metabolic output have remained incompletely defined. In this study, we demonstrate that Na^+^ deprivation induces coordinated remodeling of gut microbiome structure and function, characterized by increased alpha diversity, broad taxonomic restructuring, and reduced microbial fermentative output (Fig.1). Given that short-chain fatty acids (SCFAs) contribute approximately 5–10% of daily energy supply in humans and play central roles in appetite regulation, insulin sensitivity, and hepatic metabolism (27–30), these findings identify luminal Na^+^ availability as an under-appreciated determinant of microbiome-mediated host energy homeostasis.

Mechanistically, the microbial restructuring observed during Na^+^-deprivation is consistent with known physiological effects of luminal Na^+^ on epithelial transport and fermentation dynamics. Intraluminal Na⁺ is essential for the activity of epithelial Na^+^ transporters, including ENaC and NHE3, which regulate osmotic gradients, water flux, and substrate availability for microbial fermentation (31–33). Reduced Na^+^ availability alters luminal osmolarity and slows fermentative turnover - conditions that disproportionately disadvantage classical saccharolytic Firmicutes, particularly butyrate-producing Clostridia, whose enzymatic activity is sensitive to changes in ionic strength (34–36). This provides a biological explanation for the observed depletion of Firmicutes and the markedly reduced Firmicutes:Bacteroidetes ratio in NaD animals.

By contrast, Bacteroidetes possess extensive polysaccharide utilization loci (PULs) that confer metabolic flexibility and permit utilization of complex carbohydrates even under reduced Na^+^ or low-fermentation conditions (37, 38). Their relative enrichment in NaD animals likely reflects a competitive advantage under such environmental constraints. Similarly, the expansion of Actinobacteria and Proteobacteria is consistent with microbial responses to ionic or nutritional stress, as these taxa harbor robust stress-response regulons, siderophore biosynthesis pathways, and enhanced metabolic adaptability (39, 40). Collectively, these ecological patterns suggest that dietary Na^+^ restriction reshapes the gut microbiome by favoring osmo-adaptive and stress-tolerant taxa over classical energy-harvesting fermenters.

The functional consequences of this ecological shift were evident in both metagenomic and metabolomic profiles. Na^+^-deprivation was associated with coordinated suppression of pathways involved in carbohydrate fermentation, amino-acid metabolism, and energy metabolism, alongside enrichment of stress-response and nutrient-scavenging functions. These functional changes were mirrored by robust reductions in fecal SCFA concentrations, providing a direct metabolic readout of impaired fermentative capacity. In particular, acetate, butyrate, and hexanoate - key products of microbial carbohydrate fermentation - were markedly reduced, with butyrate showing the greatest proportional decrease (Fig 3). Because butyrate serves as the primary energy source for colonocytes, and acetate and hexanoate contribute substantially to systemic energy balance and metabolic signaling (27, 41–44), their suppression represents a physiologically meaningful consequence of Na^+^ deprivation. Prior studies have shown that lower SCFA availability reduces energy harvest and adiposity (45–47), supporting our conclusion that decreased microbial fermentation efficiency is a physiologically meaningful outcome of Na^+^ deprivation.

In addition to these major fermentative products, valerate concentrations were modestly but significantly reduced, indicating that Na^+^ restriction also influences select secondary fermentation processes. In contrast, propionate exhibited only a non-significant downward trend and branched-chain SCFAs derived predominantly from amino-acid fermentation (isobutyrate and isovalerate) were not significantly altered (Fig. 3). This pattern suggests that dietary Na^+^-deprivation preferentially impairs carbohydrate-driven microbial metabolism, rather than broadly suppressing all fermentative outputs. The selective reduction in SCFA availability provides a plausible mechanistic explanation for the reduced cecal mass and attenuated weight gain observed in Na^+^-deprived animals (Fig. 4), supporting a model in which Na^+^ restriction limits host energy harvest primarily through microbiome-mediated reductions in fermentative efficiency rather than through reduced intake or gross structural changes in the intestine. The associated host phenotypes - including reduced weight gain, decreased cecal size, and modest reductions in intake (Fig. 4) - are consistent with diminished microbial fermentation, altered luminal osmotic gradients, and delayed substrate turnover. Together, these findings establish a mechanistic link between dietary Na^+^ intake, gut microbial metabolic state, and host energy balance. They also raise an important translational question: could dietary Na^+^ restriction confer metabolic benefits that extend beyond blood-pressure reduction?

Global dietary guidelines from the World Health Organization and the American Heart Association recommend reducing Na^+^ intake to ≤2 g/day. Our findings suggest that Na^+^ reduction at this level could also influence microbial fermentation efficiency and caloric salvage, with potential implications for obesity and metabolic disease. Human studies have demonstrated that increasing propionate delivery or altering SCFA flux can reduce weight gain and improve metabolic parameters (28, 48, 49). Conversely, our data indicate that Na^+^ restriction may reduce SCFA availability and microbial energy harvest - a concept that warrants careful clinical investigation. It should be noted, however, that the relationship between dietary Na^+^, colonic pH, microbial composition, and host endocrine responses is complex. In rodents, Na^+^ deprivation induces secondary hyperaldosteronism, which stimulates apical Na⁺/H⁺ exchange in the colon (50) and may lower luminal pH sufficiently to influence microbial community structure (51, 52). Conversely, alterations in the microbiome itself can modulate aldosterone status, as increased aldosterone levels have been reported in germ-free mice and following antibiotic-mediated depletion of gut microbial biomass (53).

## CONCLUSION

In summary, this study identifies dietary Na^+^ availability as a key modulator of gut microbiome structure and metabolic function, establishing a physiological link between dietary electrolytes, microbial energy harvesting, and host metabolic outcomes. Although the interactions among Na^+^ intake, microbial diversity, luminal pH, and aldosterone signaling are complex, our findings highlight Na^+^ as an important ecological variable shaping microbial fermentative efficiency. While prolonged Na^+^ deprivation is not feasible in humans, future longitudinal studies examining stepwise Na^+^ reduction may help determine whether dietary Na^+^ modulation could complement existing strategies for managing obesity, metabolic syndrome, and gut dysbiosis.

## Availability of data and materials

The shotgun metagenomic sequencing data generated in this study have been deposited in the NCBI Sequence Read Archive (SRA) under BioProject accession [PRJNA1401895; link to the BioProject for peer-review purpose https://dataview.ncbi.nlm.nih.gov/object/PRJNA1401895?reviewer=ulodnmlnnnfer8nu8cgg514oan] (SRA: [SRR36808275 to SRR36808294]). The sample metadata required to interpret the sequencing data (diet group assignment, replicate identifiers, and phenotype measures) and the processed outputs supporting the findings of this study (taxonomic profiles, SEED Subsystems functional profiles, and fecal SCFA concentrations) are provided in the Supplementary Information Tables (ST1–ST3). Additional materials are available from the corresponding authors upon reasonable request.

## Ethics approval and consent to participate

All animal procedures were approved by the West Virginia University Institutional Animal Care and Use Committee (WVU-IACUC) and conducted in accordance with institutional and applicable national guidelines for the care and use of laboratory animals.

## Consent for publication

Not applicable.

## Authors’ contributions

Joshua Cornman-Homonoff designed the study, performed data analysis, and drafted the manuscript. Kumaran M. Rajendran was responsible for animal maintenance, collection of biological specimens, and generation of all figures. Saravanan Kolandaivelu contributed to study design and manuscript editing. Steven D. Coon and Venkatakrishna R. Jala contributed to the development of preliminary figures. Justin Kupec critically reviewed and edited the manuscript. Lei Wang and Gangqing Hu performed SRA data submission and contributed to bioinformatics figure preparation (Figures 1A–C and 2). Geoffrey I. Sandle contributed to study design and critically reviewed the manuscript. Vazhaikkurichi M. Rajendran conceived the study, supervised data analysis, secured funding, and oversaw the overall experimental design and manuscript preparation. All authors read and approved the final manuscript.

## Ethics approval and consent to participate

All animal procedures were approved by the Institutional Animal Care and Use Committee (IACUC) of West Virginia University, Morgantown, West Virginia, USA, and were conducted in accordance with institutional and national guidelines. The authors declare that they have no competing interests.

## Supplementary Figures- S1A–S1T and S2–S3

**Supplementary Figure S1.**
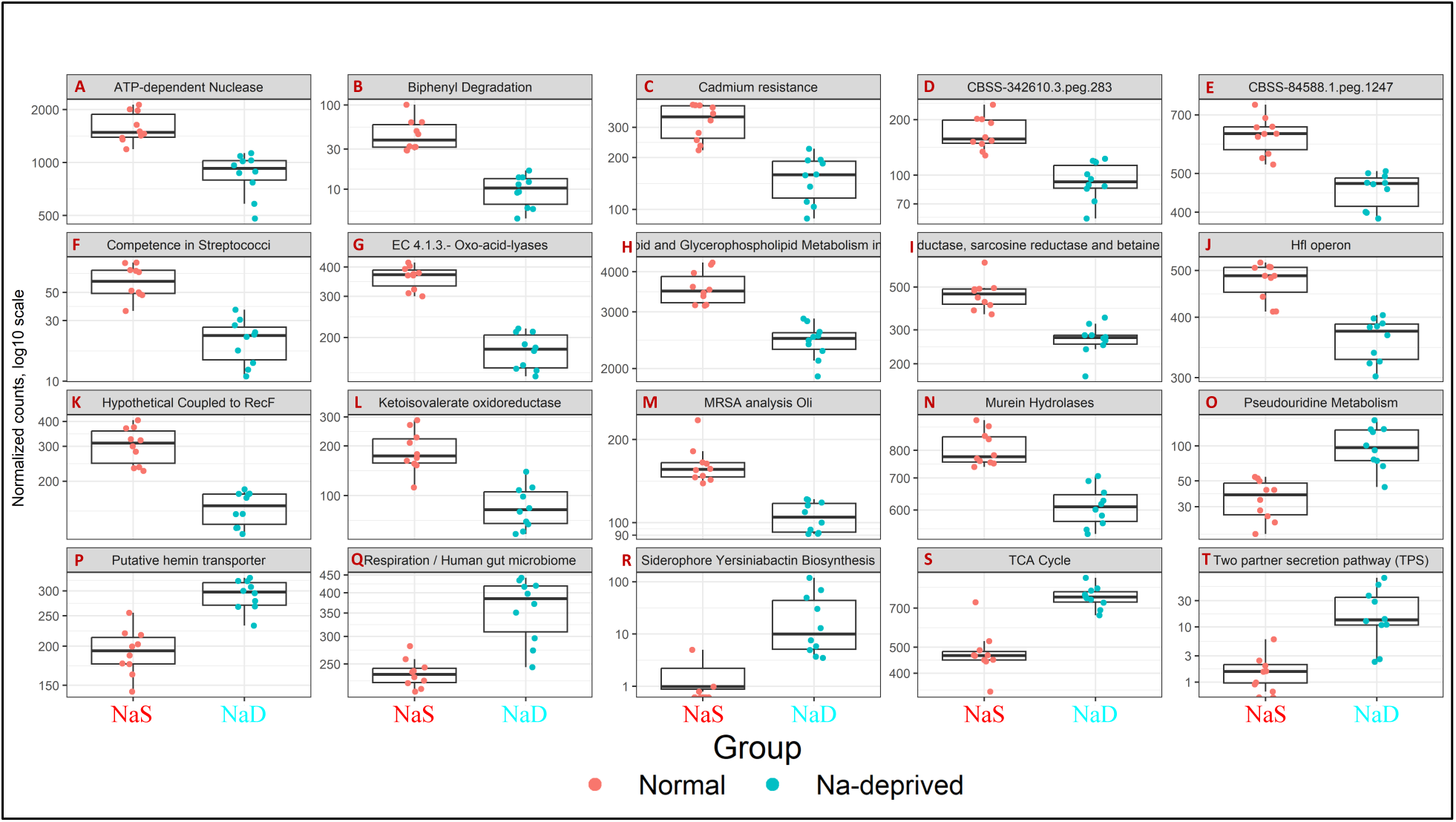
A – S1T depict relative abundances of selected functional pathways derived from shotgun metagenomic sequencing. While individual pathways did not reach statistical significance after correction for multiple testing, coordinated directional shifts in pathway representation distinguish Na^+^-deprived (NaD) and Na^+^-sufficient (NaS) microbiomes, consistent with PERMANOVA-identified differences in functional community structure.

**Supplementary Figure-S1A**: **ATP-dependent nuclease pathways**. Relative abundance of ATP-dependent nuclease–associated pathways in NaS and NaD rats. Normalized counts (log₁₀ scale) were lower in NaD microbiomes, indicating reduced representation of microbial DNA maintenance functions under Na^+^-deprivation.

**Supplementary Figure-S1B**: **Biphenyl degradation pathways**. Relative abundance of biphenyl degradation–associated pathways in NaS and NaD microbiomes. Normalized counts (log₁₀ scale) were reduced in NaD animals, suggesting decreased microbial capacity for aromatic compound metabolism during Na^+^-deprivation.

**Supplementary Figure-S1C**: **Cadmium-resistance pathways**. Relative abundance of cadmium-resistance–associated pathways in NaS and NaD rats. Normalized counts (log₁₀ scale) were decreased in NaD microbiomes, consistent with altered microbial metal-stress response potential under Na^+^-deprivation.

**Supplementary Figure-S1D**: **Stress-associated protein CBSS-342610.3.peg.283**. Relative abundance of the stress-associated protein CBSS-342610.3.peg.283 in NaS and NaD rats. Normalized counts (log₁₀ scale) were lower in NaD microbiomes, indicating altered microbial stress-response capacity under Na^+^-deprivation.

**Supplementary Figure-S1E**: **Stress-associated protein CBSS-84588.1.peg.1247**. Relative abundance of a second CBSS-associated stress protein (CBSS-84588.1.peg.1247) in NaS and NaD rats. Normalized counts (log₁₀ scale) were reduced in NaD microbiomes, consistent with coordinated suppression of microbial stress-response pathways under Na^+^-deprivation.

**Supplementary Figure-S1F**: **Streptococcal competence-associated pathways**. Competence-associated gene pathways in Streptococci were decreased in NaD rats, indicating reduced representation of functions related to horizontal gene transfer and microbial adaptability under Na^+^-deprivation.

**Supplementary Figure-S1G**: **Oxo-acid lyase pathways (EC 4.1.3)**. Relative abundance of oxo-acid lyase (EC 4.1.3)–associated pathways in NaS and NaD microbiomes. Normalized counts (log₁₀ scale) were reduced in NaD animals, consistent with altered microbial carbon metabolic potential under Na^+^-deprivation.

**Supplementary Figure-S1H**: **Lipid and membrane biosynthesis pathways**. Genes involved in lipid and membrane biosynthesis pathways were decreased in NaD microbiomes, indicating reduced representation of microbial membrane biogenesis and turnover functions under Na^+^-deprivation.

**Supplementary Figure-S1I**: **Dimethylglycine, sarcosine, and betaine-related metabolism**. Reductase-mediated pathways associated with dimethylglycine, sarcosine, and betaine metabolism were reduced under Na^+^-deprivation conditions, consistent with suppressed microbial redox and methyl-group metabolic activity.

**Supplementary Figure-S1J**: **Hfl operon (stress-response protease system)**. Relative abundance of the Hfl operon in NaS and NaD rats. Normalized counts (log₁₀ scale) were decreased in NaD microbiomes, suggesting reduced representation of stress-response proteolytic functions during Na^+^-deprivation.

**Supplementary Figure-S1K**: **RecF-associated DNA repair pathways**. RecF-associated hypothetical protein–linked pathways involved in DNA repair and replication were reduced in NaD microbiomes, consistent with diminished microbial proliferative and repair capacity under Na^+^-deprivation.

**Supplementary Figure-S1L**: **Ketoisovalerate oxidoreductase pathways**. Relative abundance of ketoisovalerate oxidoreductase–associated pathways was decreased in NaD rats, indicating reduced representation of branched-chain amino acid metabolism under Na^+^-deprivation.

**Supplementary Figure-S1M**: **MRSA-like metabolic pathways**. Genes associated with MRSA-like metabolic activity were reduced in NaD microbiomes, reflecting broader downregulation of microbial stress-handling and defense-related functions under Na^+^-deprivation.

**Supplementary Figure-S1N**: **Murein-hydrolyzing enzymes**. Relative abundance of murein-hydrolyzing enzyme pathways was decreased in NaD microbiomes, consistent with reduced representation of bacterial cell wall remodeling functions under Na^+^-deprivation.

**Supplementary Figure-S1O**: **Pseudouridine modification pathways**. Genes supporting RNA stability through pseudouridine modification were increased in NaD rats, indicating enrichment of RNA modification pathways under Na^+^-deprivation.

**Supplementary Figure-S1P**: **Hemin transporter pathways**. Hemin transporter–associated pathways were increased in NaD microbiomes, reflecting enhanced representation of microbial iron acquisition systems during Na^+^-deprivation.

**Supplementary Figure-S1Q**: **Respiration-associated pathways**. Relative abundance of respiration-associated pathways in NaS and NaD animals. Normalized counts (log₁₀ scale) were higher in NaD microbiomes, suggesting increased representation of microbial respiratory functions under Na^+^-deprivation.

**Supplementary Figure-S1R**: **Yersiniabactin siderophore biosynthesis**. Yersiniabactin siderophore biosynthesis pathways were enriched in NaD rats, indicating increased microbial iron-scavenging capacity under Na^+^-deprivation.

**Supplementary Figure-S1S**: **TCA cycle (energy metabolism)**. Genes involved in the tricarboxylic acid (TCA) cycle were enriched in NaD microbiomes, indicating increased representation of microbial oxidative energy metabolism pathways under Na^+^-deprivation.

**Supplementary Figure-S1T**: **Two-partner secretion (TPS) pathway**. Two-partner secretion (TPS) pathway genes were enriched in NaD microbiomes, reflecting increased representation of secretion-related functions during Na^+^-deprivation.

**Supplementary Figure S2.**
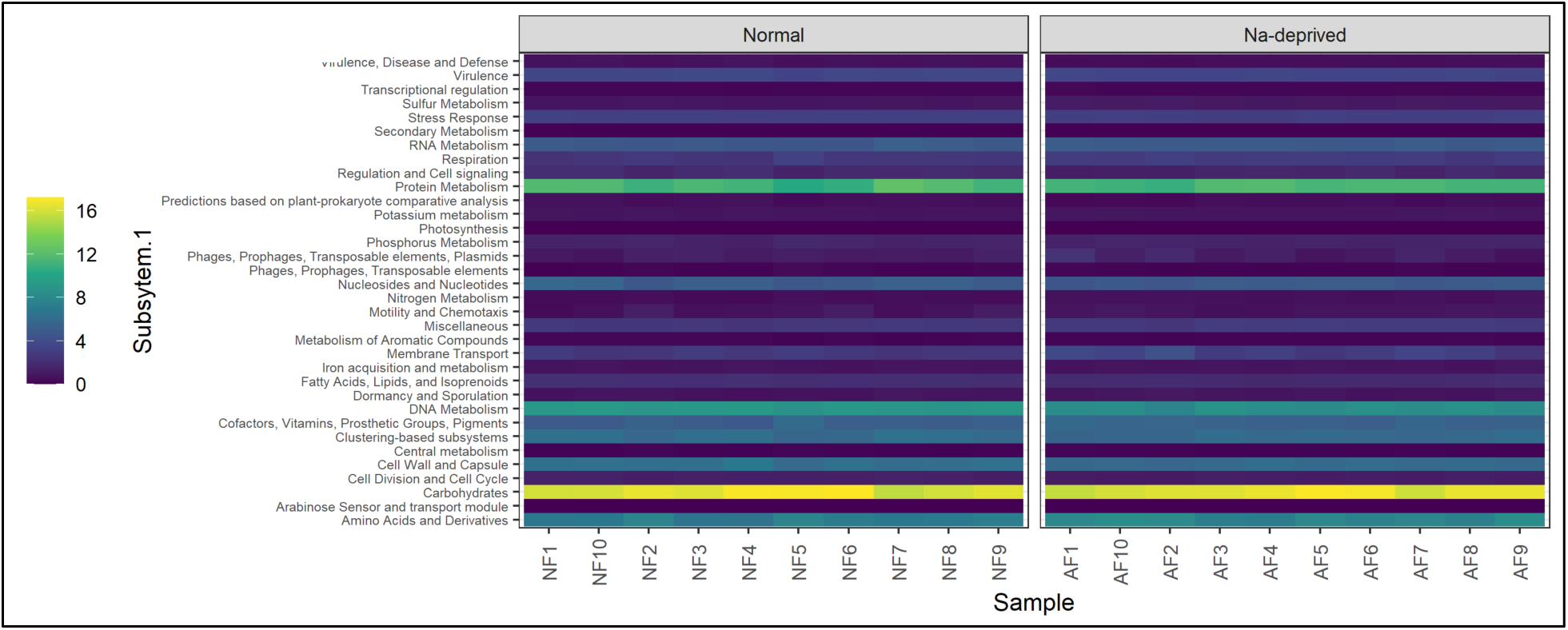
Global functional profile at SEED Subsystems Level 1. Heatmap of SEED Subsystems Level 1 functional categories derived from shotgun metagenomic sequencing of fecal samples from sodium-sufficient (NaS; Normal) and Na^+^-deprived (NaD) rats. Normalized functional abundances are shown for individual samples. This visualization provides a global overview of major functional classes and illustrates broad shifts in metabolic and stress-associated functions under Na^+^-deprived conditions.

**Supplementary Figure S3.**
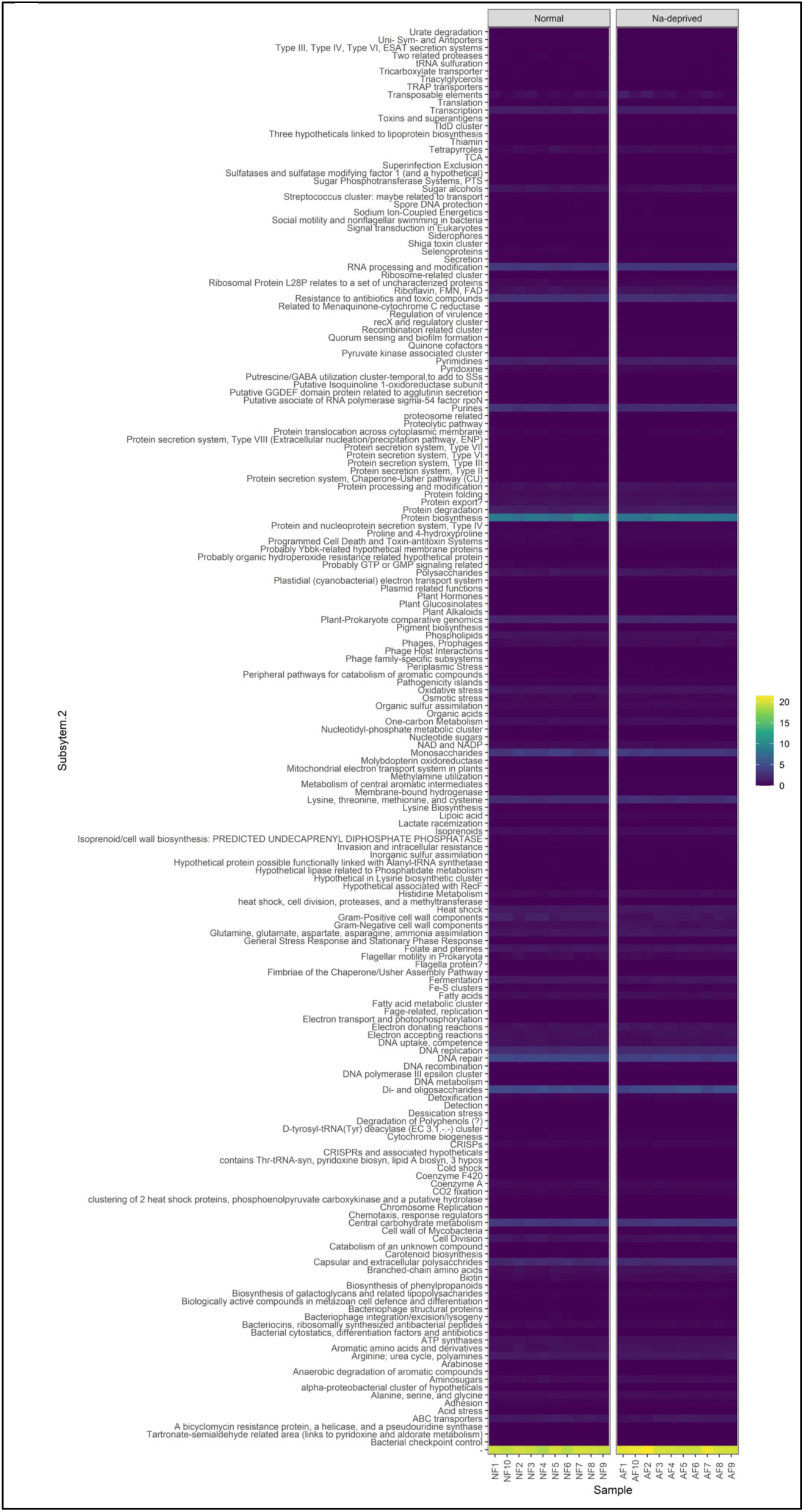
Global functional profile at SEED Subsystems Level 2. Heatmap of SEED Subsystems Level 2 functional categories derived from shotgun metagenomic sequencing of fecal samples from Na^+^-sufficient (NaS; Normal) and Na^+^-deprived (NaD) rats. Normalized functional abundances are displayed for individual samples, highlighting finer-scale functional restructuring across metabolic, stress-response, and cellular processes. Statistical inference for differentially abundant pathways is presented in the main figures and Supplementary Tables.

**Supplementary Table ST1:** Phylum-level differential abundance. Differentially abundant bacterial phyla between sodium-sufficient (NaS) and sodium-deprived (NaD) rats identified by DESeq2 analysis of shotgun metagenomic data. Log₂ fold changes, Wald test P values, and false discovery rate (FDR)–adjusted P values are shown.

|  | S00QB-0004 | S00QB-0019 | S00QB-0017 | S00QB-0009 | S00QB-0008 | S00QB-0006 | S00QB-0012 | S00QB-0011 | S00QB-0013 |
| --- | --- | --- | --- | --- | --- | --- | --- | --- | --- |
| 151781 | 90.20893803 | 67.94211348 | 47.26037969 | 79.70537675 | 82.99912136 | 78.9292322 | 53.99775952 | 45.21899655 | 63.50462 |
| 1694 | 2.244340254 | 21.29392218 | 16.78179206 | 4.097206171 | 7.007192487 | 10.94676909 | 6.76655655 | 2.935082954 | 19.94664 |
| 1117 | 0.069097955 | 0.096276957 | 0.205397319 | 0.140124812 | 0.117004852 | 0.109384636 | 0.146797633 | 0.220348748 | 0.174554 |
| 2132 | 0.061898012 | 0.0700348 | 0.203022784 | 0.105996475 | 0.113914158 | 0.066266944 | 0.119727005 | 0.195186191 | 0.172289 |
| 1298 | 0.016549121 | 0.018887667 | 0.068149168 | 0.041170692 | 0.0281106 | 0.038699896 | 0.04467754 | 0.064905288 | 0.045943 |
| 61435 | 0.016549121 | 0.017049044 | 0.057463759 | 0.047129608 | 0.032967405 | 0.04099715 | 0.037194603 | 0.06161299 | 0.045458 |
| 1005039 | 0.000322385 | 0.001002885 | 0.005223978 | 0.001805732 | 0.002354815 | 0.002297254 | 0.001980778 | 0.003762625 | 0.001132 |
| 328812 | 3.710656882 | 3.87631754 | 21.71465207 | 10.85299174 | 5.87290771 | 5.401727888 | 32.87100516 | 41.89424671 | 9.487918 |
| 281093 | 0.004943244 | 0.007020195 | 0.026357344 | 0.016251589 | 0.009860786 | 0.01431366 | 0.015406048 | 0.03221748 | 0.015692 |
| 591197 | 0.001289542 | 0.001170032 | 0.007598514 | 0.00451433 | 0.00323787 | 0.003180813 | 0.004621815 | 0.007054923 | 0.003235 |
| 1457365 | 0.000644771 | 0.000835737 | 0.002137082 | 0.002166879 | 0.001471759 | 0.001236983 | 0.001320518 | 0.002351641 | 0.001617 |
| 833 | 0.000859695 | 0.001504327 | 0.003086896 | 0.003611464 | 0.003532222 | 0.00282739 | 0.004621815 | 0.004938446 | 0.002911 |
| 861299 | 0.000644771 | 0.001671475 | 0.005936339 | 0.003069745 | 0.00323787 | 0.00194383 | 0.004401728 | 0.007290087 | 0.00388 |
| 456827 | 0.000537309 | 0.000501442 | 0.000949814 | 0.001083439 | 0.000147176 | 0.001236983 | 0 | 0.000940656 | 0.000323 |
| 239935 | 1.515534144 | 1.065899571 | 3.700476094 | 0.242690396 | 0.039590321 | 0.150558498 | 0.159782731 | 1.18969511 | 1.441088 |
| 406548 | 0.026328148 | 0.033596646 | 0.102105026 | 0.072590431 | 0.0482737 | 0.058668335 | 0.063164798 | 0.112878761 | 0.074416 |
| 389348 | 0.001934313 | 0.002172917 | 0.003799257 | 0.002708598 | 0.002501991 | 0.003180813 | 0.002641037 | 0.00517361 | 0.002426 |
| 2697043 | 0.001397004 | 0.001170032 | 0.006411246 | 0.002166879 | 0.001177407 | 0.002650678 | 0.00242095 | 0.005643938 | 0.004691 |
| 1930593 | 0.000537309 | 0.000501442 | 0.000237454 | 0.001264012 | 0.000883055 | 0.000706847 | 0.001100432 | 0.000705492 | 0.000161 |
| 543 | 1.213781335 | 4.370572681 | 6.35568167 | 2.761325552 | 2.188358679 | 2.446398878 | 3.629665006 | 5.090596964 | 3.102028 |
| 848 | 0.069635264 | 0.091596827 | 0.170254194 | 0.115025136 | 0.100373974 | 0.071745012 | 0.114004758 | 0.180841182 | 0.15077 |
| 157 | 0.054375684 | 0.08140083 | 0.167167298 | 0.134888189 | 0.110823464 | 0.124405144 | 0.1181864 | 0.194480699 | 0.128772 |
| 248039 | 0.028692308 | 0.062346015 | 0.810903867 | 0.048574194 | 0.01766111 | 0.199684393 | 0.320445807 | 0.058791021 | 0.027178 |
| 1197717 | 0.013432728 | 0.015043274 | 0.083346195 | 0.030878019 | 0.018396989 | 0.030217727 | 0.032572788 | 0.052911919 | 0.038017 |
| 2108365 | 0.011605877 | 0.021394879 | 0.035855486 | 0.022029932 | 0.020015924 | 0.015020507 | 0.025750109 | 0.039272402 | 0.025236 |
| 392734 | 0.008596946 | 0.012368915 | 0.035618032 | 0.024016237 | 0.021340508 | 0.023149253 | 0.018707345 | 0.041153715 | 0.021839 |
| 2899133 | 0.002256698 | 0.000835737 | 0.002611989 | 0.002528025 | 0.002649166 | 0.001590407 | 0.002200864 | 0.005643938 | 0.000485 |
| 2735562 | 0.000322385 | 0.001002885 | 0.000474907 | 0.000722293 | 0.001030231 | 0.000176712 | 0.000440173 | 0.000940656 | 0.000970 |
| 28262 | 0.003223855 | 0.00200577 | 0.010922863 | 0.008848087 | 0.007064444 | 0.006361627 | 0.007923111 | 0.014109845 | 0.006147 |
| 64160 | 0.002256698 | 0.003175802 | 0.004986525 | 0.00505605 | 0.004120926 | 0.003534237 | 0.004621815 | 0.005879102 | 0.004853 |
| 171695 | 0.00236416 | 0.00468013 | 0.006886153 | 0.006861782 | 0.006181388 | 0.005301356 | 0.007482938 | 0.006114266 | 0.006794 |
| 1408281 | 0.001289542 | 0.001002885 | 0.004274164 | 0.003611464 | 0.001618935 | 0.003887661 | 0.003301296 | 0.007054923 | 0.003073 |
| 14 | 0.00118208 | 0.001671475 | 0.004511617 | 0.003611464 | 0.001618935 | 0.001413695 | 0.002641037 | 0.005408774 | 0.00210 |
| 2847778 | 0.000967156 | 0 | 0.000237454 | 0.00054172 | 0.000147176 | 0.000706847 | 0.000880346 | 0.00117582 |  |
| 187145 | 0.000967156 | 0.00133718 | 0.004749071 | 0.002528025 | 0.001471759 | 0.001767119 | 0.000660259 | 0.002586805 | 0.001941 |
| 936456 | 0.000752233 | 0.000167147 | 0.002374535 | 0.001444586 | 0.001471759 | 0.001236983 | 0.001980778 | 0.002586805 | 0.001617 |
| 693075 | 0 | 0.000334295 | 0.001424721 | 0.001083439 | 0 | 0.000530136 | 0.000880346 | 0.000470328 | 0.000485 |

**Supplementary Table ST2:**
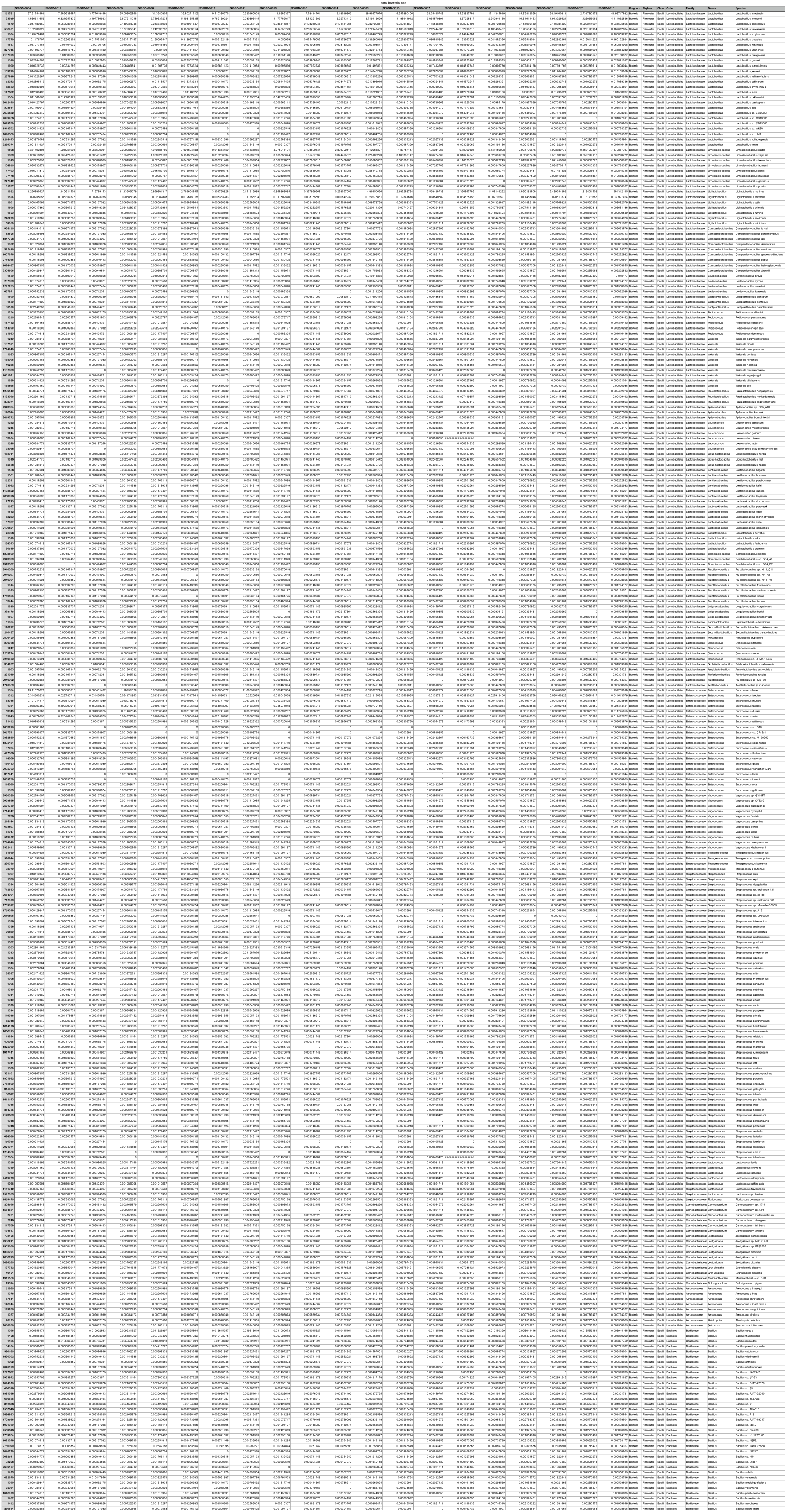
Species-level differential abundance: Differentially abundant bacterial species between sodium-sufficient (NaS) and sodium-deprived (NaD) rats identified by DESeq2 analysis of shotgun metagenomic data. Log₂ fold changes, Wald test P values, and FDR-adjusted P values are reported.

**Supplementary Table ST3:** Fecal short-chain fatty acid concentrations: Fecal short-chain fatty acid concentrations in sodium-sufficient (NaS) and sodium-deprived (NaD) rats. Values represent normalized concentrations measured by gas chromatography. Group comparisons were performed using two-sided Mann–Whitney U tests, as described in the Methods.

|  |  |  |  | scfa_concentration_r |
| --- | --- | --- | --- | --- |
| MBI.Sample.ID | Acetic.Acid.mmol.per.kg | Propionic.Acid.mmol.per.kg | Isobutyric.Acid.mmol.per.kg | Butyric.Acid.mmol.per.kg |
| S00QB-0001 | 40.23707181 | 5.962095339 | 0.314071986 |  |
| S00QB-0002 | 53.16740291 | 7.296917202 | 1.085531794 |  |
| S00QB-0003 | 38.96901263 | 7.325940661 | 1.063297526 |  |
| S00QB-0004 | 60.44804821 | 10.70584856 | 1.208309259 |  |
| S00QB-0005 | 52.76419684 | 7.336504056 | 1.053369976 |  |
| S00QB-0006 | 57.00986081 | 7.437522321 | 0.586609337 |  |
| S00QB-0007 | 59.1270533 | 7.658327872 | 0.450455128 |  |
| S00QB-0008 | 59.54463204 | 7.980166929 | 1.013728519 |  |
| S00QB-0009 | 37.97820666 | 6.36679418 | 1.189727203 |  |
| S00QB-0010 | 58.27817336 | 8.559927393 | 0.398978546 |  |
| S00QB-0011 | 16.05191387 | 3.000355327 | 0.794017208 |  |
| S00QB-0012 | 38.5933428 | 6.931126048 | 0.991608379 |  |
| S00QB-0013 | 47.49552327 | 6.666348933 | 0.671752731 |  |
| S00QB-0014 | 29.94133053 | 5.005230689 | 0.65943853 |  |
| S00QB-0015 | 36.18635733 | 6.056835523 | 0.581644685 |  |
| S00QB-0016 | 21.79993436 | 5.289698213 | 0.561529655 |  |
| S00QB-0017 | 58.99376908 | 12.22392944 | 0.211266717 |  |
| S00QB-0018 | 34.29018763 | 5.697464471 | 0.709920813 |  |
| S00QB-0019 | 22.71066116 | 3.787072684 | 0.78793842 |  |
| S00QB-0020 | 20.41361833 | 3.912540111 | 0 |  |

## REFERENCES

1. Thursby E, Juge N. Introduction to the human gut microbiota. Biochem J. 2017;474(11):1823–36.

2. Arumugam M, Raes J, Pelletier E, Le Paslier D, Yamada T, Mende DR, et al. Enterotypes of the human gut microbiome. Nature. 2011;473(7346):174–80.

3. Flint HJ, Duncan SH, Scott KP, Louis P. Interactions and competition within the microbial community of the human colon: links between diet and health. Environ Microbiol. 2007;9(5):1101–11.

4. Walker AW, Ince J, Duncan SH, Webster LM, Holtrop G, Ze X, et al. Dominant and diet-responsive groups of bacteria within the human colonic microbiota. ISME J. 2011;5(2):220–30.

5. Walter J, Ley R. The human gut microbiome: ecology and recent evolutionary changes. Annu Rev Microbiol. 2011;65:411–29.

6. Postler TS, Ghosh S. Understanding the Holobiont: How Microbial Metabolites Affect Human Health and Shape the Immune System. Cell Metab. 2017;26(1):110–30.

7. Schroeder BO, Backhed F. Signals from the gut microbiota to distant organs in physiology and disease. Nat Med. 2016;22(10):1079–89.

8. Singh RK, Chang HW, Yan D, Lee KM, Ucmak D, Wong K, et al. Influence of diet on the gut microbiome and implications for human health. J Transl Med. 2017;15(1):73.

9. Donaldson GP, Lee SM, Mazmanian SK. Gut biogeography of the bacterial microbiota. Nat Rev Microbiol. 2016;14(1):20–32.

10. Pagliai G, Dinu M, Fiorillo C, Becatti M, Turroni S, Emmi G, et al. Modulation of gut microbiota through nutritional interventions in Behcet’s syndrome patients (the MAMBA study): study protocol for a randomized controlled trial. Trials. 2020;21(1):511.

11. Rinninella E, Tohumcu E, Raoul P, Fiorani M, Cintoni M, Mele MC, et al. The role of diet in shaping human gut microbiota. Best Pract Res Clin Gastroenterol. 2023;62-63:101828.

12. Ayten S, Bilici S. Modulation of Gut Microbiota Through Dietary Intervention in Neuroinflammation and Alzheimer’s and Parkinson’s Diseases. Curr Nutr Rep. 2024;13(2):82–96.

13. Dasriya VL, Samtiya M, Ranveer S, Dhillon HS, Devi N, Sharma V, et al. Modulation of gut-microbiota through probiotics and dietary interventions to improve host health. J Sci Food Agric. 2024;104(11):6359–75.

14. Hindle VK, Veasley NM, Holscher HD. Microbiota-Focused Dietary Approaches to Support Health: A Systematic Review. J Nutr. 2025;155(2):381–401.

15. Juraschek SP, Miller ER, 3rd, Weaver CM, Appel LJ. Effects of Sodium Reduction and the DASH Diet in Relation to Baseline Blood Pressure. J Am Coll Cardiol. 2017;70(23):2841–8.

16. Gupta DK, Lewis CE, Varady KA, Su YR, Madhur MS, Lackland DT, et al. Effect of Dietary Sodium on Blood Pressure: A Crossover Trial. JAMA. 2023;330(23):2258–66.

17. Sacks FM, Svetkey LP, Vollmer WM, Appel LJ, Bray GA, Harsha D, et al. Effects on blood pressure of reduced dietary sodium and the Dietary Approaches to Stop Hypertension (DASH) diet. DASH-Sodium Collaborative Research Group. N Engl J Med. 2001;344(1):3–10.

18. Pimenta E, Gaddam KK, Oparil S, Aban I, Husain S, Dell’Italia LJ, et al. Effects of dietary sodium reduction on blood pressure in subjects with resistant hypertension: results from a randomized trial. Hypertension. 2009;54(3):475–81.

19. Adnan S, Nelson JW, Ajami NJ, Venna VR, Petrosino JF, Bryan RM, Jr., et al. Alterations in the gut microbiota can elicit hypertension in rats. Physiol Genomics. 2017;49(2):96–104.

20. Durgan DJ, Ganesh BP, Cope JL, Ajami NJ, Phillips SC, Petrosino JF, et al. Role of the Gut Microbiome in Obstructive Sleep Apnea-Induced Hypertension. Hypertension. 2016;67(2):469–74.

21. Li J, Zhao F, Wang Y, Chen J, Tao J, Tian G, et al. Gut microbiota dysbiosis contributes to the development of hypertension. Microbiome. 2017;5(1):14.

22. Smiljanec K, Lennon SL. Sodium, hypertension, and the gut: does the gut microbiota go salty? Am J Physiol Heart Circ Physiol. 2019;317(6):H1173–H82.

23. Naqvi S, Asar TO, Kumar V, Al-Abbasi FA, Alhayyani S, Kamal MA, et al. A cross-talk between gut microbiome, salt and hypertension. Biomed Pharmacother. 2021;134:111156.

24. Yan X, Jin J, Su X, Yin X, Gao J, Wang X, et al. Intestinal Flora Modulates Blood Pressure by Regulating the Synthesis of Intestinal-Derived Corticosterone in High Salt-Induced Hypertension. Circ Res. 2020;126(7):839–53.

25. Wu C, Yosef N, Thalhamer T, Zhu C, Xiao S, Kishi Y, et al. Induction of pathogenic TH17 cells by inducible salt-sensing kinase SGK1. Nature. 2013;496(7446):513–7.

26. Nagase S, Karashima S, Tsujiguchi H, Tsuboi H, Miyagi S, Kometani M, et al. Impact of Gut Microbiome on Hypertensive Patients With Low-Salt Intake: Shika Study Results. Front Med (Lausanne). 2020;7:475.

27. Byrne CS, Chambers ES, Morrison DJ, Frost G. The role of short chain fatty acids in appetite regulation and energy homeostasis. Int J Obes (Lond). 2015;39(9):1331–8.

28. Chambers ES, Viardot A, Psichas A, Morrison DJ, Murphy KG, Zac-Varghese SE, et al. Effects of targeted delivery of propionate to the human colon on appetite regulation, body weight maintenance and adiposity in overweight adults. Gut. 2015;64(11):1744–54.

29. Frampton J, Murphy KG, Frost G, Chambers ES. Short-chain fatty acids as potential regulators of skeletal muscle metabolism and function. Nat Metab. 2020;2(9):840–8.

30. Rajendran VM, Nanda Kumar NS, Tse CM, Binder HJ. Na-H Exchanger Isoform-2 (NHE2) Mediates Butyrate-dependent Na+ Absorption in Dextran Sulfate Sodium (DSS)-induced Colitis. J Biol Chem. 2015;290(42):25487–96.

31. Hooper LV, Macpherson AJ. Immune adaptations that maintain homeostasis with the intestinal microbiota. Nat Rev Immunol. 2010;10(3):159–69.

32. Kunzelmann K, Mall M. Electrolyte transport in the mammalian colon: mechanisms and implications for disease. Physiol Rev. 2002;82(1):245–89.

33. Louis P, Flint HJ. Formation of propionate and butyrate by the human colonic microbiota. Environ Microbiol. 2017;19(1):29–41.

34. Duncan SH, Belenguer A, Holtrop G, Johnstone AM, Flint HJ, Lobley GE. Reduced dietary intake of carbohydrates by obese subjects results in decreased concentrations of butyrate and butyrate-producing bacteria in feces. Appl Environ Microbiol. 2007;73(4):1073–8.

35. Macfarlane GT, Macfarlane S. Fermentation in the human large intestine: its physiologic consequences and the potential contribution of prebiotics. J Clin Gastroenterol. 2011;45 Suppl:S120–7.

36. Tuncil YE, Xiao Y, Porter NT, Reuhs BL, Martens EC, Hamaker BR. Reciprocal Prioritization to Dietary Glycans by Gut Bacteria in a Competitive Environment Promotes Stable Coexistence. mBio. 2017;8(5).

37. Martens EC, Chiang HC, Gordon JI. Mucosal glycan foraging enhances fitness and transmission of a saccharolytic human gut bacterial symbiont. Cell Host Microbe. 2008;4(5):447–57.

38. Zhao G, Nyman M, Jonsson JA. Rapid determination of short-chain fatty acids in colonic contents and faeces of humans and rats by acidified water-extraction and direct-injection gas chromatography. Biomed Chromatogr. 2006;20(8):674–82.

39. Merchant SS, Helmann JD. Elemental economy: microbial strategies for optimizing growth in the face of nutrient limitation. Adv Microb Physiol. 2012;60:91–210.

40. Song Y, Wu X, Li Z, Ma QQ, Bao R. Molecular mechanism of siderophore regulation by the Pseudomonas aeruginosa BfmRS two-component system in response to osmotic stress. Commun Biol. 2024;7(1):295.

41. Koh A, De Vadder F, Kovatcheva-Datchary P, Backhed F. From Dietary Fiber to Host Physiology: Short-Chain Fatty Acids as Key Bacterial Metabolites. Cell. 2016;165(6):1332–45.

42. He J, Zhang P, Shen L, Niu L, Tan Y, Chen L, et al. Short-Chain Fatty Acids and Their Association with Signalling Pathways in Inflammation, Glucose and Lipid Metabolism. Int J Mol Sci. 2020;21(17).

43. Facchin S, Bertin L, Bonazzi E, Lorenzon G, De Barba C, Barberio B, et al. Short-Chain Fatty Acids and Human Health: From Metabolic Pathways to Current Therapeutic Implications. Life (Basel). 2024;14(5).

44. O’Riordan KJ, Collins MK, Moloney GM, Knox EG, Aburto MR, Fulling C, et al. Short chain fatty acids: Microbial metabolites for gut-brain axis signalling. Mol Cell Endocrinol. 2022;546:111572.

45. Rahat-Rozenbloom S, Fernandes J, Gloor GB, Wolever TM. Evidence for greater production of colonic short-chain fatty acids in overweight than lean humans. Int J Obes (Lond). 2014;38(12):1525–31.

46. Cuevas-Sierra A, Ramos-Lopez O, Riezu-Boj JI, Milagro FI, Martinez JA. Diet, Gut Microbiota, and Obesity: Links with Host Genetics and Epigenetics and Potential Applications. Adv Nutr. 2019;10(suppl_1):S17–S30.

47. May KS, den Hartigh LJ. Modulation of Adipocyte Metabolism by Microbial Short-Chain Fatty Acids. Nutrients. 2021;13(10).

48. Lu B, Hanyaloglu AC, Ma Y, Frampton AE, Limb C, Merali N, et al. Propionate Induces Energy Expenditure via Browning in Mesenteric Adipose Tissue. J Clin Endocrinol Metab. 2025;111(1):256–67.

49. Blaak EE, Canfora EE, Theis S, Frost G, Groen AK, Mithieux G, et al. Short chain fatty acids in human gut and metabolic health. Benef Microbes. 2020;11(5):411–55.

50. Cho JH, Musch MW, Bookstein CM, McSwine RL, Rabenau K, Chang EB. Aldosterone stimulates intestinal Na+ absorption in rats by increasing NHE3 expression of the proximal colon. Am J Physiol. 1998;274(3):C586–94.

51. Duncan SH, Louis P, Thomson JM, Flint HJ. The role of pH in determining the species composition of the human colonic microbiota. Environ Microbiol. 2009;11(8):2112–22.

52. Xie Z, He W, Gobbi A, Bertram HC, Nielsen DS. The effect of in vitro simulated colonic pH gradients on microbial activity and metabolite production using common prebiotics as substrates. BMC Microbiol. 2024;24(1):83.

53. Moore BN, Medcalf AD, Muir RQ, Xu C, Marques FZ, Pluznick JL. Commensal microbiota regulate aldosterone. Am J Physiol Renal Physiol. 2024;326(6):F1032–F8.

